# DiaThor: R package for computing diatom metrics and biotic indices

**DOI:** 10.1101/2021.03.31.437895

**Authors:** M.M. Nicolosi Gelis, M.B. Sathicq, J. Jupke, J. Cochero

**Author notes:** **Corresponding author:** M.M. Nicolosi Gelis.

## Abstract

1. Diatoms are widely used to detect changes in water quality due to their specific sensibility to a variety of environmental conditions. Among the different diatom-based tools to measure water quality, the biotic indices, ecological and morphological traits are the most commonly used.
2. *DiaThor*, contains 18 functions that provide morphological data of the samples (number and shape of chloroplasts, total biovolume), ecological data (species richness, evenness, diversity, size classes, ecological guilds, ecological preferences) and biotic indices (Descy Index, EPID Index, Indice Diatomique Artois-Picardie, Swiss Diatom Index, Pampean Diatom Index, ILM index, Specific Pollution sensitivity Index, Lobo Index, Sládecek index, SPEAR_herbicides_ index and the Trophic index). A web application (in Shiny) was also developed to provide access to the package for those users not familiar with the R environment.
3. The structure of the package, how it functions and the structure of the input and output data are explained. To demonstrate the most common use of *DiaThor*, an example of the package performance is also provided.
4. The *DiaThor* package aims to contribute to the water quality assessment based on diatom assemblages, while also providing researchers with an open platform to suggest new statistics and functionalities to be integrated into future builds.

## Introduction

The continuing need for new and improved methods for monitoring the environmental quality and the changes in aquatic ecosystems, leads to the creation of different indices and metrics based on physical, chemical and biological parameters. One of the most utilized groups used to monitor freshwater ecosystems are diatoms. Diatoms have been used to assess river water quality since the early work by Kolkwitz and Marsson (1908), while the first environmental quality indices based on diatoms assemblages were developed about 60 years ago (*e.g.* Zelinka & Marvan, 1961). Nowadays, diatoms are part of routine bioassessment protocols (Stevenson et al., 2010), for example as one of four biological quality elements in the European Water Framework Directive.

This group exhibits an important diversity and their community composition is strongly structured by numerous environmental factors (Patrick, 1961; Lange-Bertalot, 1979) and responds rapidly to changes in nutrient concentrations even in environments where the internal loading is not a problem (Kelly & Whitton, 1995; Nicolosi Gelis, 2020).

Numerous studies dealing with water quality assessment have focused on the application of standardized methodologies based on diatom assemblages, using taxonomic diversity as an important tool for biomonitoring, since it allows ecological assessment even under subtle environmental impacts (Gomez & Licursi, 2001; Birks, 2010).

Diatom-based environmental assessment traditionally required specific taxonomic knowledge, but there is only a little loss of ecological information when the taxonomic accuracy decreased from the species level to higher taxonomic groups (Rimet & Bouchez, 2012, B-Bères et al., 2014). Thus, the application of diatom functional groups (life-forms and ecological guilds), have become increasingly popular in ecological assessments over the past decade (Tapolczai et al., 2016; Riato et al., 2017). Functional diversity metrics can improve our knowledge of community and ecosystem responses to environmental changes at different spatial and temporal scales (Péru & Dolédec, 2010) and the role by which important environmental drivers influence diatom community structure (Lange et al., 2016).

Depending on the objective of the analysis, on the environment, the spatial scale of the study and on the kind of disturbance, there are different choices of indices or ecological features to be used for biomonitoring. The calculation of biotic indices and the classification by ecological and/or morphological traits are time consuming and prone to errors, particularly when several sites need to be monitored. Software has been developed to assist the researchers in these tasks. The most widely used solution is OMNIDIA (Lecointe et al., 1993), a proprietary software that assists in diatom counting, diatom inventory management, and the calculation of diatom indices for water quality assessment.

Here we present DiaThor, a free and open source R package for calculating biotic and ecological indices and metrics for diatom samples, such as morphological and ecological traits. To provide access to the package for those users not familiar with the R environment, a web application was also developed using the package Shiny.

## Description

***DiaThor*** contains 19 functions in total (Table 1). The main functions are *diat_loadData()* and *diaThorAll().*

**Figure 1.**
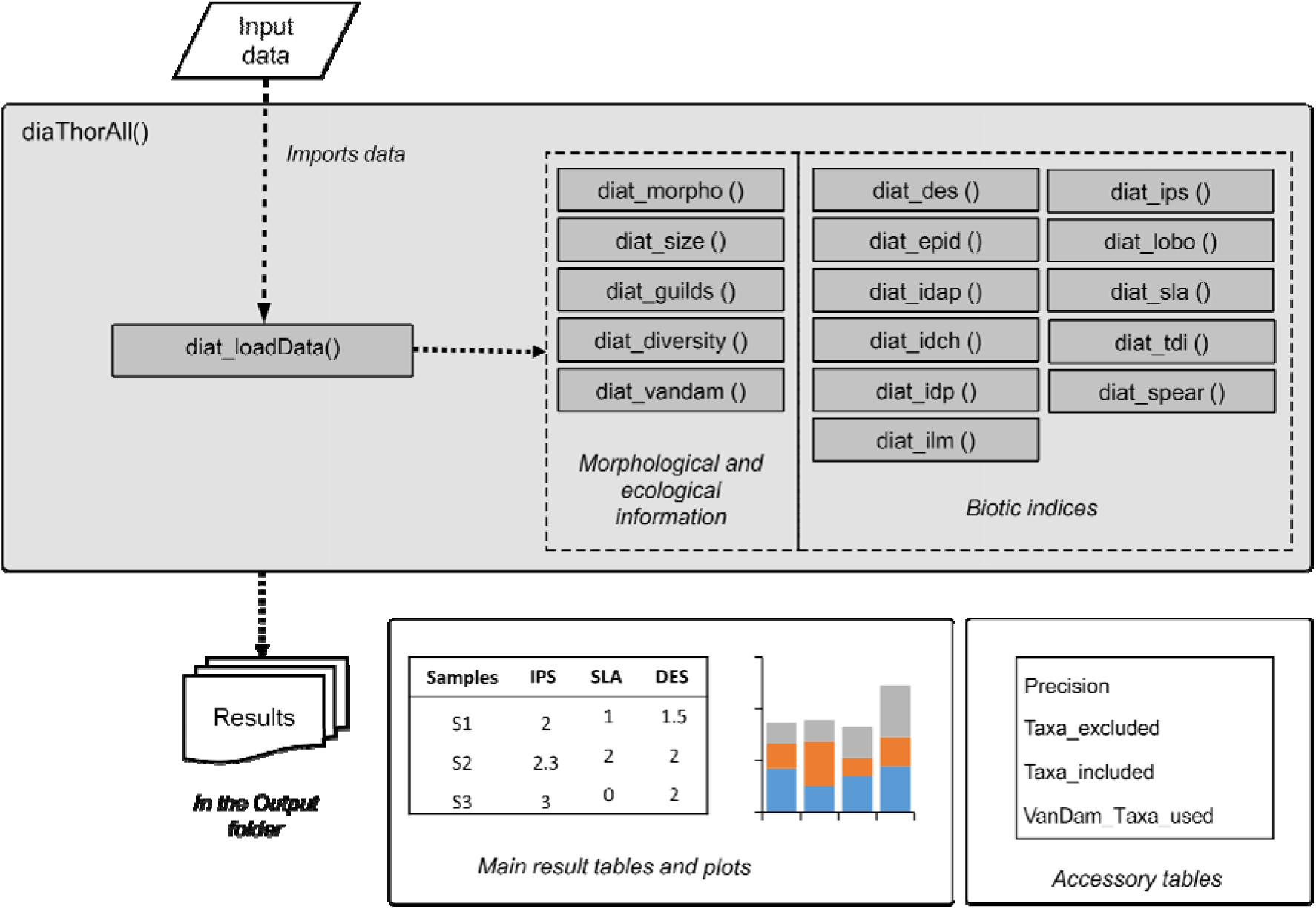
Workflow of the diaThorAll() function, which includes all other individual functions of the package DiaThor to output tables and plots.

The *diat_loadData()* function checks and converts the input data into the format needed by the package. It loads the input data (a CSV file or an R data.frame) and sets the output folder for the results. The other functions in the package use the resulting list from the *diat_loadData()* function function to match the input taxa names with their value for each metric. To do this, it first attempts an exact match; if it fails for a taxon (e.g. misspelled taxa), it uses a heuristic search to find a similar name that differs from the input data by a defined number of characters set in the *maxDistTaxa* argument of the function (default: 2). Consider that the higher the *maxDistTaxa* value is, the higher are the chances that a taxon is incorrectly mistaken for another, that multiple taxa are matched to the same taxon, and the longer the algorithm will take to find a result. However, decreasing the value lowers the probability of finding an accronym. (a *maxDistTaxa* value of zero is equivalent to running an exact match). The main function is the *diaThorAll()* function (Figure 1). With eleven optional arguments (described in Table S1), this function runs all other functions included in the package (Table 1) one after the other, producing all possible outputs, and exports them to the Output folder.

The other 16 functions allow the user to calculate each ecological or biotic index separately, and can be run individually using the output produced by the *diat_loadData()* as the argument *resultLoad*.

## Indices calculated

In its current version ***DiaThor*** calculates morphological data of the samples, ecological data, and biotic indices.

The **morphological data** provided for each sample includes: (i) the proportion of cells with a given number of chloroplasts (e.g. 1 chloroplast, 3 chloroplasts, 2-rarely-4 chloroplasts, etc.); (ii) the proportion of cells with each shape of chloroplast (e.g. H-shaped chloroplast, lobed, etc.); and (iii) the total biovolume of the sample.

The **ecological data** calculated for each sample includes: (i) species richness, evenness and diversity by using the *vegan* package (Oksanen et al., 2015); (ii) the proportion of size classes of diatoms (as defined by Rimet & Bouchez, 2012); (iii) the proportion of each ecological guild (i.e. planktonic, low profile, high profile and motile, as defined by Rimet & Bouchez, 2012); and (iv) the ecological indicator indices defined by Van Dam (1994), which include salinity, N-uptake metabolism, oxygen requirements, saprobity, trophic state and moisture.

For these calculations, the script retrieves the Diat.Barcode database (Rimet et al., 2019) by using the *diatbarcode* package (Keck 2020), to obtain information for each diatom species. Factor levels that are not used are filtered out (i.e. if no species with lobed chloroplasts are present in the sample, that percentage of cells will not be included in the output).

The **biotic indices** calculated for each sample are among the most commonly used for evaluating the water quality in freshwater environments: Descy Index (Descy, 1979), EPID index (Dell’Uomo, 1996), Indice Diatomique Artois-Picardie (Prygiel & Coste, 1993), Swiss Diatom Index (Hürlimann & Niederhaus, 2007), Pampean Diatom Index (Gómez & Licursi, 2001), ILM index (Leclercq & Maquet, 1987), Specific Pollution Sensitivity Index (IPS; Coste 1982), Lobo Index (Lobo et al. 2002, 2004), Sládecek index (Sládecek, 1986), SPEAR_herbicides_ index (Wood et al., 2019) and the Trophic index (Kelly et al., 1995).

## Input data and data structure

The input file for the package is an R data.frame or an external comma-separated value (CSV) file. If a data.frame is not specified as an argument of the *diat_loadData()* function, a dialog box will be shown so the user can select the CSV file manually.

Diatom taxa should be listed as rows. The first row has to contain the headers with the sample names. The taxa names have to be included in column 1 (not as row names), and that column has to be labeled “species”. If the input data contains a column named “acronym”, that column will be automatically used to match taxa with their ecological values. This is more accurate than the heuristic search of species’ names that the *diat_findAcronym()* function will perform.

The taxa names do not have to include the taxa authors. The format to input taxa names is: Generic name + specific epithet + subspecies (if exists, indicated by “subsp.”) + variety (if exists, indicated by “var.”) + form (if exists, indicated by “f.”).

For example, valid names include: *“Achnanthes hungarica”, “Rhopalodia gibba* var. *gibba”, “Achnanthes lanceolata* subsp. *frequentissima”*.

The other columns (samples) have to contain either the absolute or relative abundance of each species in each sample.

The performance of the package is highly dependent on the proper naming of the species in the input data. It is recommended to input the taxa names along with their standard acronyms; the *diat_findAcronym* function can be run previously to help this process and check that all taxa are being correctly recognized by the script.

The *diat_loadData()* generates an R list with five objects that can be used to calculate the biotic indices: (i) *taxaIn* and (ii) *taxaInRA* are dataframes containing the taxa abundances and relative abundances respectively; (iii) the vectors *sampleNames* contains the names of the samples as stated in the input file; (iv) the *resultsPath* vector contains the path for the Output folder; and (v) *taxaInEco* is a dataframe resulting from joining the input data with the Diat.Barcode database. Generally, there is no need for the user to modify this resulting R list, which has to be entered as an argument into the other functions of the package to obtain the results.

## Internal databases and additional packages needed

The package contains internally multiple databases (in .rda format): (i) for each biotic index calculated (TDI, IPS, IDP, etc.) it includes a table with the taxa name recognized by that index, and the necessary ecological information to calculate the index; (ii) sample data, to test the package as described in the “Example” section, and; (iii) an offline copy of the database provided openly by the Diat.Barcode project (Rimet et al., 2019), in case the user does not have access to the internet (current version: 9.0). If connected to the internet, the Diat.Barcode database will be updated automatically through *diatbarcode* package (Rimet et al., 2021). ***DiaThor*** depends on other packages to properly function, which are installed when the library is loaded: *stringdist* (van der Loo, 2014), *vegan* (Oskanen et al. 2020), *permute* (Simpson, 2019), *lattice* (Deepayan, 2008), *ggplot2* (Wickham, 2016), *purrr* (Henry & Wickham, 2020), *data.table* (Dowle & Srinivasan, 2020) and *tidyr* (Wickham, 2020).

## Outputs

The outputs of the package include multiple files (Table S2). The *DiaThor_results.csv* file contains all the biotic and ecological indices calculated per sample. For each of the ecological and morphological variables calculated, a column is also added specifying the percentage of abundance that did not fall within the classified categories (columns end in “*Indet*”) and the number of taxa used per sample and index (columns end in *“Taxa.used”*).*The plots.pdf* file shows lollipop figures and barchart plots for each index (sample in Figure 2).

**Figure 2.**
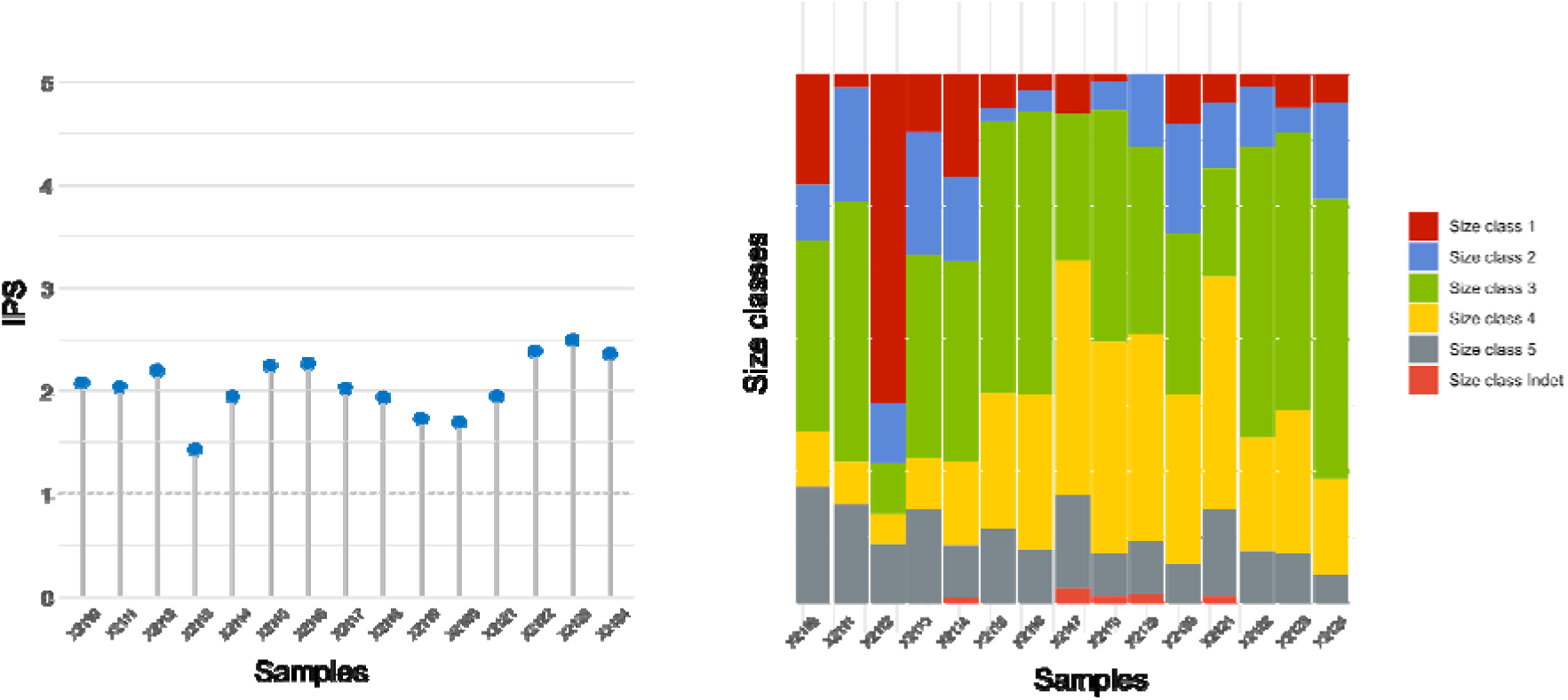
Example of the outputs obtained for biotic indices (IPS, left) and ecological traits (size classes, right).

The other files exported provide the user with additional information to improve their calculations, and they are important to be checked before reporting the data. The *Precision.csv* file shows what percentage of taxa were actually used for the calculation of each index (not all indices include ecological information for all existing taxa). The *Recognized_acronyms.csv* file will show the user which taxa were identified and their acronyms found, and will list the updated taxon name and acronym if the taxonomy has been updated. The *Taxa_included.csv* and the *Taxa_excluded.csv* files will list which taxa were actually included or excluded from the calculation of each biotic index. Finally, the *VanDam_Taxa_used.txt* file lists the taxa included for the calculation of each ecological index following VanDam’s classification, whenever they had a relative abundance in the sample > 0.

**Table 1.** Functions included in DiaThor and their description.

| Function name | Description |
| --- | --- |
| *diaThorAll* | The main function, includes all the other functions |
| *diat_loadData* | Loads the data into DiaThor in the correct format |
| *diat_morpho* | Calculate morphological parameters for diatoms (Rimet et al., 2019) |
| *diat_guilds* | Calculates the proportion of ecological guilds for diatoms (Rimet et al., 2012) |
| *diat_diversity* | Calculates species' richness and the Shannon diversity index (Shannon & Weaver, 1949) |
| *diat_vandam* | Calculates ecological information for diatoms based on the Van Dam classification (Van Dam, 1994) |
| *diat_size* | Calculate size classes for diatoms (Rimet et al., 2012) |
| *diat_des* | Calculates the Descy Index (DES, Descy, 1979) |
| *diat_epid* | Calculates the EPID index (EPID; Dell'Uomo, 1996) |
| *diat_idap* | Calculates the Indice Diatomique Artois-Picardie (IDAP; Prygiel & Coste, 1993) |
| *diat_idch* | Calculates the Swiss Diatom Index (IDCH; Hürlimann & Niederhauser, 2007) |
| *diat_idp* | Calculates the Pampean Diatom Index (IDP; Gómez & Licursi, 2001) |
| *diat_ilm* | Calculates the ILM Index (ILM; Leclercq & Maquet, 1987) |
| *diat_ips* | Calculates the Specific Pollutosensitivity Index (IPS; Coste, 1982) |
| *diat_lobo* | Calculates the Lobo Index (LOBO; Lobo et al., 2002, 2004) |
| *diat_sla* | Calculates the Sládecek Index (SLA; Sládecek, 1986) |
| *diat_spear* | Calculates the SPEAR(herbicides) Index (SPEAR; Wood et al., 2019) |
| *diat_tdi* | Calculates the Trophic index (TDI; Kelly et al., 1995) |

## Example

To demonstrate the most common use of DiaThor, the package includes sample data with the abundance of 164 diatom species in 108 sampled sites (Nicolosi Gelis et al., 2020).

~~~
# Installing the package and load it into the R environment
> *install. packages(“diathor”)*
> *library(diathor)*
# Load the internally included sample data
> *data(“diat_sampleData”)*
# Run diaThorAll to get all the outputs from the sample data with the default settings, and store the results into the “results” object, to also retain the output within R
> *results* <- *diaThorAll(diat_sampleData)*
# The package will request an Output folder through a dialog box
*[1] “Select Results folder”*
~~~

After the Results folder is selected, all the calculations conducted will be shown in the console.

## Implementation and availability

The package is available for the R programming language, and free to be downloaded from the CRAN repository. A Shiny web app has also been developed to provide access online to the package for those users not familiar with the R language, and it can be found at: https://limnolab.shinyapps.io/diathor-shiny/

More information about the package can be found at https://cran.r-project.org/web/packages/diathor/index.html. It can be installed directly from the R environment running *install.packages*(“*diathor*”), and this manuscript describes version 0.0.5 of the software.

The source code for the package is available at https://github.com/limnolab/DiaThor, and the source code for the Shiny web app at https://github.com/limnolab/DiaThor-Shiny. Both codes are open under an MIT License. Any issues can be reported through the GitHub Issues system, and collaborative development is encouraged. Each output is calculated in a separate R script within the package, allowing the incorporation of other indices in future releases.

## Conclusions

We introduced the open-source R package *DiaThor*, which is designed to calculate diatom indexes and morphological and ecological data. The current version of the *DiaThor* package contains functions to calculate 11 biotic indexes and 5 ecological and morphological classifications.

In the package repository, we provide the source code for both, the package and the Shiny web app. The package will be updated frequently to include new functionalities, and is completely open to encourage users to contribute to the project adding their own indices or improving the current ones. We hope that *DiaThor* will become an important tool for dealing with water quality assessment based on diatom assemblages.

## Acknowledgements

The authors would like to thank Dr. Rebecca Wood for her valuable help in testing *DiaThor* and improving the calculation of the SPEAR(herbicide) index.

## Author contributions

**Conceptualization**: M.M.N.G., M.B.S., J.C.; **Data curation:** M.M.N.G., M.B.S., J.C.; **Methodology**: M.M.N.G., M.B.S.; **Project administration**: M.M.N.G., M.B.S., J.C.; **Software**: M.B.S., M.M.N.G., J.C., J.K.; **Supervision**: J.C.; **Validation**: M.B.S., M.M.N.G., J.C., J.K.; **Writing-original draft**: M.B.S., M.M.N.G., J.C., J.K.; **Writing-review & editing**: M.B.S., M.M.N.G., J.C., J.K.

